# Plasma and ovarian metabolomic responses to chronic stress in female mice

**DOI:** 10.1101/2022.01.03.474852

**Authors:** Oana A. Zeleznik, Tianyi Huang, Chirag J. Patel, Elizabeth M. Poole, Clary B. Clish, Guillermo N. Armaiz-Pena, Archana S. Nagaraja, A. Heather Eliassen, Katherine H. Shutta, Raji Balasubramanian, Laura D. Kubzansky, Susan E. Hankinson, Anil K. Sood, Shelley S. Tworoger

**Affiliations:** Channing Division of Network Medicine, Brigham and Women’s Hospital and Harvard Medical School, Boston, MA, USA; Departments of Nutrition and Epidemiology, Harvard T.H. Chan School of Public Health, Boston, MA, USA; Department of Biomedical Informatics, Harvard Medical School, Boston, MA, United States; Bluebird Bio, Cambridge, MA, USA; Broad Institute of the Massachusetts Institute of Technology and Harvard University, Cambridge, MA, USA; Department of Basic Sciences, Division of Pharmacology, School of Medicine, Ponce Health Sciences University, PR United States; Department of Gynecologic Oncology & Reproductive Medicine, UT MD Anderson Cancer Center, Houston, TX, USA; Department of Biostatistics and Epidemiology, School of Public Health and Health Sciences, University of Massachusetts, Amherst, MA, USA; Department of Social and Behavioral Sciences, Harvard T.H. Chan School of Public Health, Boston, MA, USA; Department of Cancer Epidemiology, Moffitt Cancer Center, Tampa, FL, USA

**Keywords:** restraint stress, chronic stress, female mice, metabolomics, plasma, ovarian tissue

## Abstract

**Background:** Chronic stress may affect metabolism of amino acids, lipids, and other small molecule metabolites, but these alterations may differ depending on tissue evaluated. We examined metabolomic changes in plasma and ovarian tissue samples from female mice due to chronic stress exposure.

**Methods:** At 12 weeks old, healthy, female, C57 black mice were randomly assigned to three weeks of chronic stress using daily restraint (2 hours/day; n=9) or normal care (n=10). Metabolomic profiling was conducted on plasma and ovarian tissues. Using the Wilcoxon Rank Test, Metabolite Set Enrichment Analysis, and Differential Network Analysis we identified metabolomic alterations occurring in response to restraint stress. All p-values were corrected for multiple testing using the false discovery rate approach.

**Results:** In plasma, individual lysophosphatidylcholines (positively) and the metabolite classes carnitines (positively), diacylglycerols and triacylglycerols (inversely) were associated with restraint stress (adjusted-p’s<0.2). In contrast, diacylglycerols and triacylglycerols were increased while carnitines were decreased in ovarian tissue from stressed mice (adjusted-p’s<0.2). However, several metabolites (cholesteryl esters, phosphatidylcholines/ phosphatidylethanolamines plasmalogens and multiple amino acids) were consistently inversely associated with restraint stress in plasma and ovarian tissue (adjusted-p’s<0.2).

**Conclusion:** We identified differences in multiple lipid and amino acid metabolites in plasma and ovarian tissue of female mice after exposure to chronic stress. Some affected metabolites (primarily triacylglycerols and diacylglycerols) exhibited opposite associations with chronic stress in plasma (a marker of systemic influences) versus in ovarian tissue (representing local changes), suggesting research to understand the biological impact of chronic stress needs to consider both systemic and tissue-specific alterations.

## Introduction

Chronic stress is a common, complex, and multifactorial process that may have long-term health impacts at least in part through prolonged neuroendocrine dysregulation (1). Sustained activation of the sympathetic nervous system and hypothalamic-pituitary-adrenal axes can induce dysregulation of stress response systems and alterations in behaviors and homeostasis, leading to systemic metabolic perturbations (2, 3) Increasing evidence suggests that chronic stress may affect metabolism of amino acids, lipids, and other small molecule metabolites in animals (4-18) and humans (19-31). For example, several studies observed that tyrosine levels were lower in individuals with a major depressive disorder (MDD) or a Type D personality (a form of psychological distress resulting from negative affectivity and social inhibition), as well as in chronically stressed versus control mice (6, 8, 20, 21, 25). Similarly, phosphatidylcholines (PCs) and certain fatty acids such as octadecanoic and hexadecanoic acids were lower in individuals with MDD/Type D personality and stressed mice (5, 10, 15, 21, 25, 28).

Moreover, there are known sex differences in stress response, with women exhibiting increased susceptibility to the adverse health consequences of chronic stress compared to men (32, 33). Additionally, there is compelling evidence for the role of chronic stress in some female-specific diseases, such as ovarian cancer, a disease with metabolic risk factors such as body size and circulating levels of specific metabolites (34-37). Specifically, chronic stress in mice leads to larger, more aggressive ovarian tumors than in controls via up-regulation of norepinephrine and decreases in dopamine levels (38-40). In parallel, mounting evidence in humans suggests that some markers for chronic stress (e.g., depression, anxiety, post-traumatic stress disorder [PTSD]) may be associated with increased risk of developing ovarian cancer (41-44).

However, the exact mechanisms through which metabolic dysregulation induced by chronic stress promotes ovarian cancer growth and progression are not fully understood. A major barrier is that nearly all animal studies were conducted on male mice or female mice seeded with ovarian cancer cells. To our knowledge, there has been only one study in healthy female mice (16), reporting that four days of restraint stress were associated with higher HDL levels and altered hepatic gene expression related to amino acid and lipid metabolism. Further, no studies have examined metabolomic changes due to chronic stress in ovarian tissues specifically, which may differ from the patterns observed in circulation. Therefore, to provide further insight into stress-induced metabolic alterations at the systemic (plasma) versus local (ovarian tissue) level, we used liquid chromatography (LC)-mass spectrometry (MS/MS) to evaluate the metabolomic dysregulation due to chronic stress in plasma and ovarian tissue samples from female mice.

## Methods and Materials

### Animal experiments

Female, immunocompetent mice (C57BL6; 10-to 12-weeks old) were obtained from Taconic Farms (Hudson, NY). All experiments were approved by the Institutional Animal Care and Use Committee of the M.D. Anderson Cancer Center. A total of 19 animals were randomly assigned to control (N=10) or chronic stress conditions (N=9) using a well-characterized restraint system, two hours/day for 21 consecutive days. This intermittent immobilization protocol has been shown to increase catecholamines in a manner characteristic of chronic stress (38); prior work has also demonstrated food intake (45) and subsequent body weight and cardiac:body weight ratio (38) to not differ between restraint-stressed and control groups. On day 22, blood was collected into a Heparin tube through the tail vein and processed within 1 hour of collection. Plasma was aliquoted and frozen at −80°C until shipped to the assay laboratory. Necropsy ensued and ovarian tissue was removed and flash frozen in 30-40 mg pieces (one sample had 5mg of tissue) until shipment.

### Metabolomic profiling

Metabolomics were measured in plasma and ovarian tissues samples using similar methods. Aqueous homogenates of ovarian tissue samples were created by weighing the tissue and adding 4uL of water per mg of tissue and then using a bead beater (TissueLyser II, QIAGEN; Germantown, MD) to homogenize the sample. Profiles of endogenous polar metabolites and lipids were measured using LC-MS/MS. Details on sample preparation and chromatography methods were similar to those described previously(46) but the MS analyses were carried out using high-resolution accurate mass (HRAM) profiling in these experiments. Briefly, for polar metabolites, hydrophilic interaction liquid chromatography (HILIC) analyses were conducted in the positive ion mode using a Nexera X2 U-HPLC (Shimadzu, Marlborough, MA)-Q Exactive orbitrap (Thermo Fisher Scientific; Waltham, MA) LC-MS instrument (HILIC-positive ion mode).

Plasma and tissue extracts (10µL) were diluted using 90 µL of 74.9:24.9:0.2 v/v/v acetonitrile/methanol/formic acid containing stable isotope-labeled internal standards (valine-d8, Isotec; and phenylalanine-d8, Cambridge Isotope Laboratories; Andover, MA). The samples were centrifuged (10 min, 9,000 x g, 4°C) and the supernatants were injected directly onto a 150 × 2 mm Atlantis HILIC column (Waters; Milford, MA). The column was eluted isocratically at a flow rate of 250 µL/min with 5% mobile phase A (10 mM ammonium formate and 0.1% formic acid in water) for 1 min followed by a linear gradient to 40% mobile phase B (acetonitrile with 0.1% formic acid) over 10 min. The electrospray ionization voltage was 3.5 kV and data were acquired using full scan analysis over m/z 70-800 at 70,000 resolution.

For lipid metabolites, assays were performed using a Nexera X2 U-HPLC (Shimadzu, Marlborough, MA) coupled to an Exactive Plus orbitrap mass spectrometer (Thermo Fisher Scientific; Waltham, MA) in the positive ion mode (C8 chromatography-positive ion mode MS). Lipids were extracted from plasma or tissue homogenates (10 µL) using 190 µL of isopropanol containing 1,2-dilauroyl-sn-glycero-3-phosphocholine as an internal standard (Avanti Polar Lipids; Alabaster, AL). After centrifugation (10 min, 9,000 x g, ambient temperature), supernatants (2 µL) were injected directly onto a 100 × 2.1 mm ACQUITY BEH C8 column (1.7 µm; Waters; Milford, MA). The column was initially eluted isocratically at a flow rate of 450 µL/min with 80% mobile phase A (95:5:0.1 vol/vol/vol 10mM ammonium acetate/methanol/formic acid) for 1 minute followed by a linear gradient to 80% mobile-phase B (99.9:0.1 vol/vol methanol/formic acid) over 2 minutes, a linear gradient to 100% mobile phase B over 7 minutes, then 3 minutes at 100% mobile-phase B. MS analyses were carried out using electrospray ionization in the positive ion mode (source voltage was 3kV, source temperature was 300°C, sheath gas was 50, auxiliary gas was 15) using full scan analysis over m/z 200-1100 and at 70,000 resolution and 3 Hz data acquisition rate.

Prior to the assays, LC-MS system sensitivity and chromatography quality were checked by analyzing reference samples: synthetic mixtures of reference metabolites (Sigma; St. Louis, MO) and a lipid extract prepared from a pooled human plasma (BioreclamationIVT; Chestertown, MD). During the application of each method, internal standard peak areas were monitored for quality control. Moreover, reference pooled plasma samples were included in each set of analyses, with samples inserted before and after the study samples. TraceFinder 3.1 software (Thermo Fisher Scientific; Waltham, MA) was used for automated peak integration and metabolite peaks were manually reviewed for quality of integration and compared against a known standard to confirm identity. Metabolites with a signal-to-noise ratio <10 were considered unquantifiable. Metabolite signals were retained as measured LC-MS peak areas, proportional to metabolite concentrations, and appropriate for metabolite clustering and correlative analyses.

A total of 268 metabolites were consistently measured across the 19 plasma samples. After excluding four metabolites with missing data in more than two of the stressed or control mice, the analysis of the plasma metabolomics data included 264 metabolites. In the 19 ovarian tissue samples, 274 metabolites were measured. We excluded one metabolite with missing data in more than two of the stressed or control mice, leaving 273 metabolites available for analysis. An overlapping set of 247 metabolites were measurable in both plasma and ovarian tissue samples.

### Statistical analysis

All analyses were conducted in plasma and ovarian tissue separately. Spearman correlations were used to present the correlation structure among metabolites. For each identified metabolite peak from plasma or ovarian tissue samples, we assessed the difference in peak area comparing mice exposed to chronic restraint stress versus controls using the Wilcoxon rank test. We calculated the percent difference between stressed and control mice as the difference between mean metabolite value among stressed mice and mean metabolite values among control mice divided by the mean metabolite value among control mice. For plotting purposes, we log-transformed and standardized (mean = 0, standard deviation =1) metabolite values.

Metabolite set enrichment analysis (MSEA) was used to identify groups of metabolites associated with restraint stress (47). We used principal component analysis to summarize metabolite classes (all principal components needed to explain 75% of the variance) which we investigated further using differential network analysis (DINGO) (48) to identify differences in partial correlations networks between restraint-stressed mice and controls. Nodes of the network reflect individual metabolite classes and an edge between a pair of nodes reflects a partial correlation that is significantly different in the stressed vs control animals. The DINGO algorithm was run in plasma and ovarian tissue samples separately. The positive FDR approach (q-value) was used to account for testing multiple correlated hypotheses in individual metabolite analyses and the assessment of significantly different partial correlations between stressed and control animals in DINGO (49). MSEA uses FDR to correct for testing multiple hypotheses. Due to the hypothesis-generating nature of the study, we report and discuss findings with q-value<0.2 (individual metabolites, DINGO) or p-adjusted<0.2 (MSEA).

## Results

### Metabolites associated with restraint stress in plasma

Restraint-stressed and control mice demonstrated different pairwise correlation structures for measured plasma metabolites (Figure 1, Panels A and B). When comparing levels of individual metabolites between restraint-stressed mice and control mice, 87 metabolites were statistically significantly different at q-value≤0.2 and 27 metabolites at q-value≤0.05 (Figure 2A, Supplementary Table 1). The top five metabolites with the lowest q-values (q-value<0.005) included four lysophosphatidylcholines (LPC; C18:2, C18:1, C20:4, C22:6) and one triacylglycerol (TAG; C58:8). The average percent difference between restraint-stressed and control mice ranged from −26% for C18:1 LPC to −47% for C58:8 TAG. In addition, levels of several amino acids, including carnosine, putrescine, thyroxine, citrulline and anserine, were decreased in restraint-stressed compared with control mice (q-value<0.1). Regarding the magnitude of the difference, exposure to restraint stress resulted in the largest decrease in C58:6 triglyceride (TAG) (65% lower; q-value=0.04) and the largest increase in C36:1 phosphatidylcholine (PC) plasmalogen (104% higher; q-value=0.03). Using MSEA, we identified three metabolite classes that were enriched at p-adjusted<0.2 (Figure 2B, Supplementary Table 2). TAGs and diacylglycerol (DAG) had a negative normalized enrichment score (NES; p-adjusted<0.03), suggesting lower plasma metabolite levels in restraint-stressed mice compared with control mice, while carnitines had a positive NES (p-adjusted<0.001) suggesting higher plasma metabolite levels in restraint-stressed mice compared with control mice.

**Figure 1.**
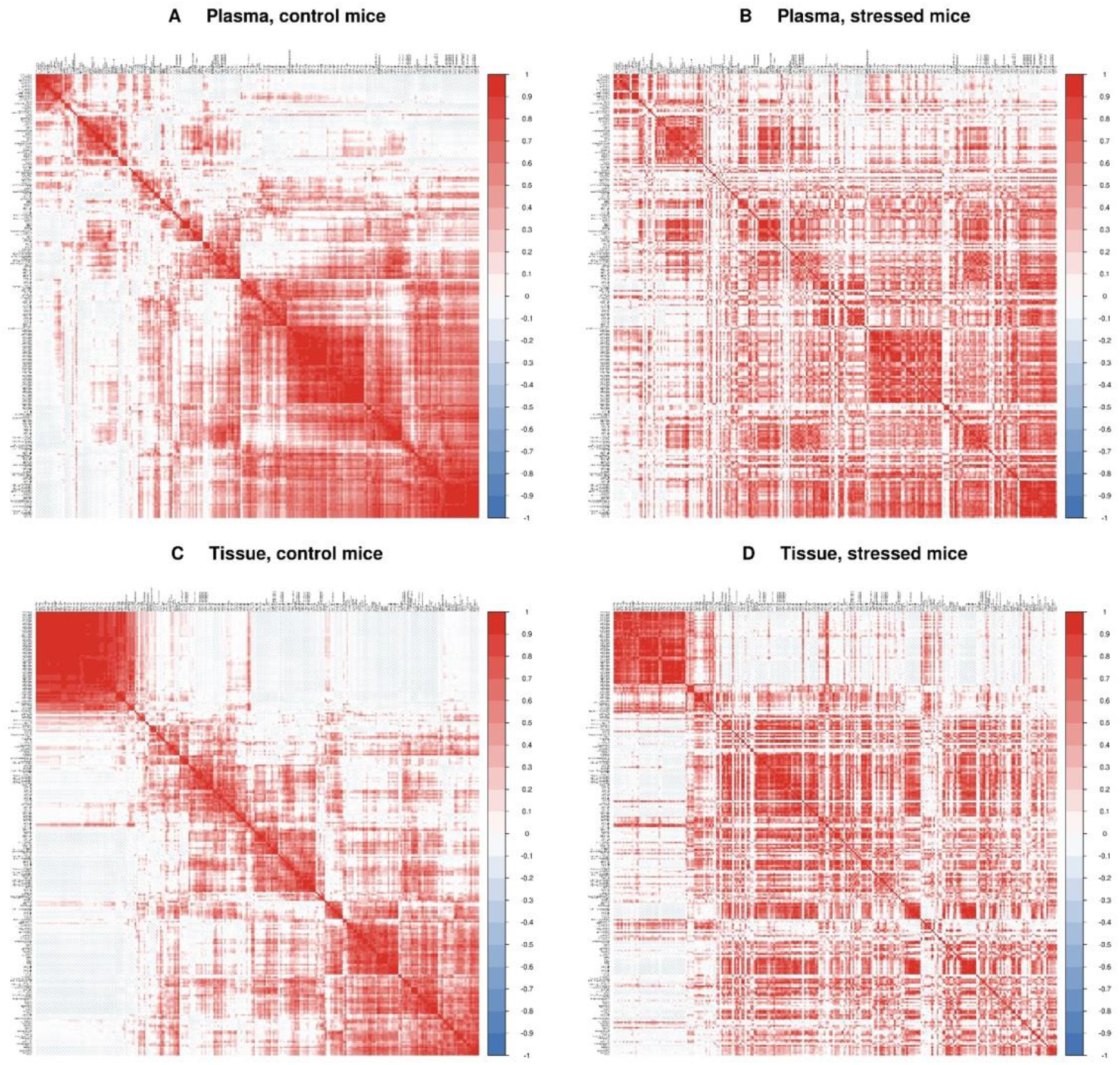
Correlation structure among metabolites. Correlations between metabolites measured in healthy female mice: A. Plasma among control mice, B. Plasma among restraint-stressed mice, C. Ovarian tissue among control mice, and D. Ovarian tissue among restraint-stressed mice. Positive correlations are shown in shades of blues while negative correlations are shown in shades of red. The order of metabolites is the same in panels A and B and in panels C and D.

**Figure 2.**
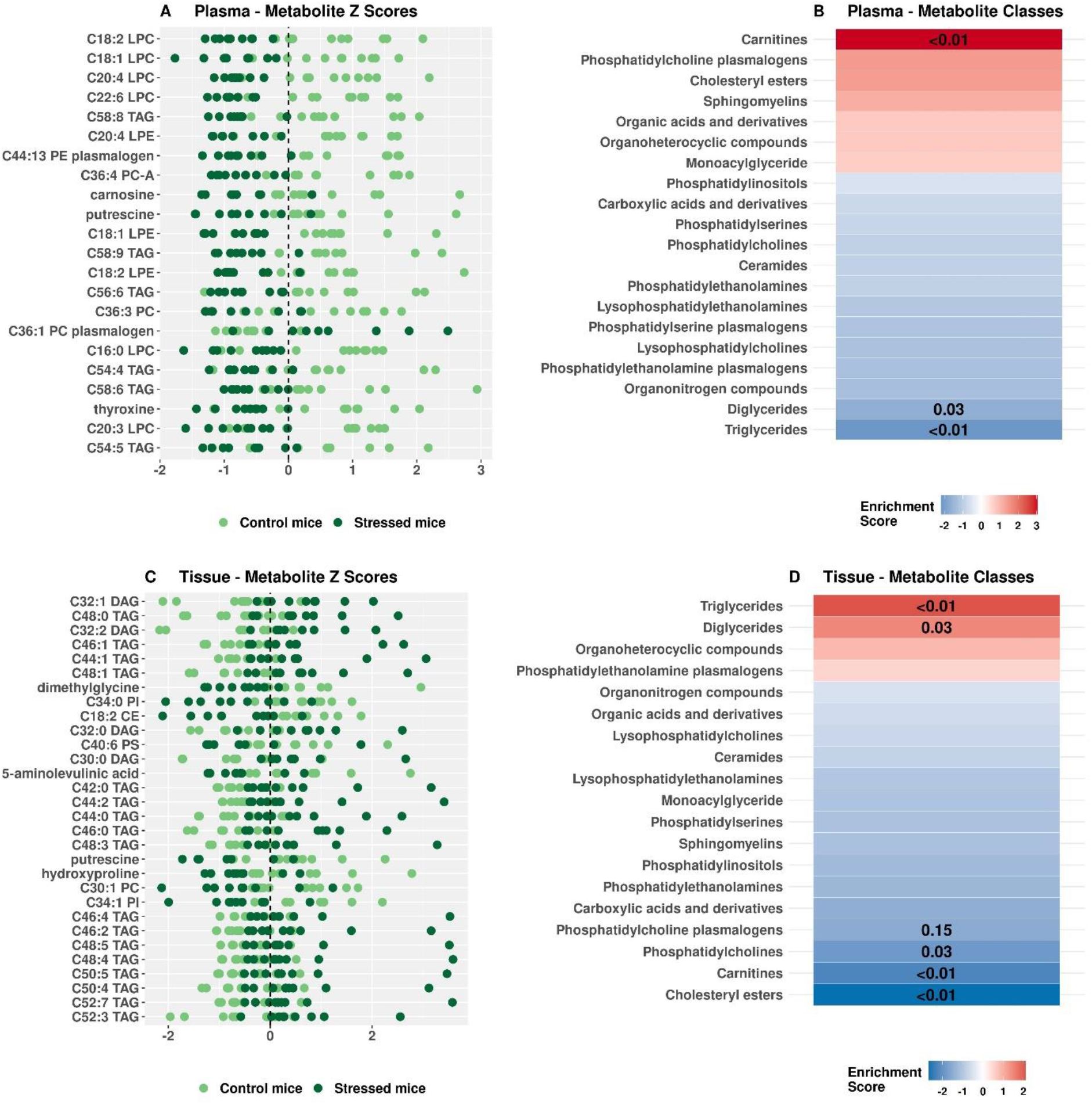
Individual metabolites and metabolite classes associated with restraint-stress in plasma and in tissue. Panels A and C: Z-scores for individual metabolites that differed between restraint-stressed and control mice at q-values≤0.05 in plasma and ovarian tissue, respectively. Each dot represents a sample. Control mice are shown in light green and restraint-stressed mice are shown in dark green. Metabolites are ordered from high to low levels in the stressed mice. **Panel B and D:** Metabolite classes enriched in restraint-stressed compared to control mice in plasma and ovarian tissue, respectively. Positive enrichment is shown in shades of red while negative enrichment is shown in shades of blue. Metabolites classes are ordered from up-regulated to down-regulated in stressed mice compared to control mice. Adjusted p-values<0.20 are overlaid on the plot.

The plasma differential network reflecting differences in partial correlation, comparing restraint-stressed and control mice included seven metabolite classes: LPC, PC plasmalogens, lysophosphatidylethanolamine (LPE) plasmalogens, phosphatidylserines (PS), ceramides, carboxylic acids and derivatives, and organic acids and derivatives (Figure 3A, Supplementary Table 3). These seven classes were connected by four statistically significant edges (q-value<0.2), reflecting differences in the pairwise partial correlations among these metabolite classes occurring in the restraint-stressed mice versus control mice. Each pair of metabolite classes is linked by two edges, one representing partial correlations between the two metabolite classes among controls animals and the other among stressed animals. Specifically, the inverse correlations between LPC and carboxylic acids and derivatives, between PC plasmalogens and organic acids and derivatives, and between PS and organic acids and derivatives were consistently stronger (more negative) in restraint-stressed mice than control mice. PE plasmalogens had a positively partial correlation with ceramides in control mice (r=0.44), but there was no correlation in restraint-stressed mice (r≈0).

**Figure 3.**
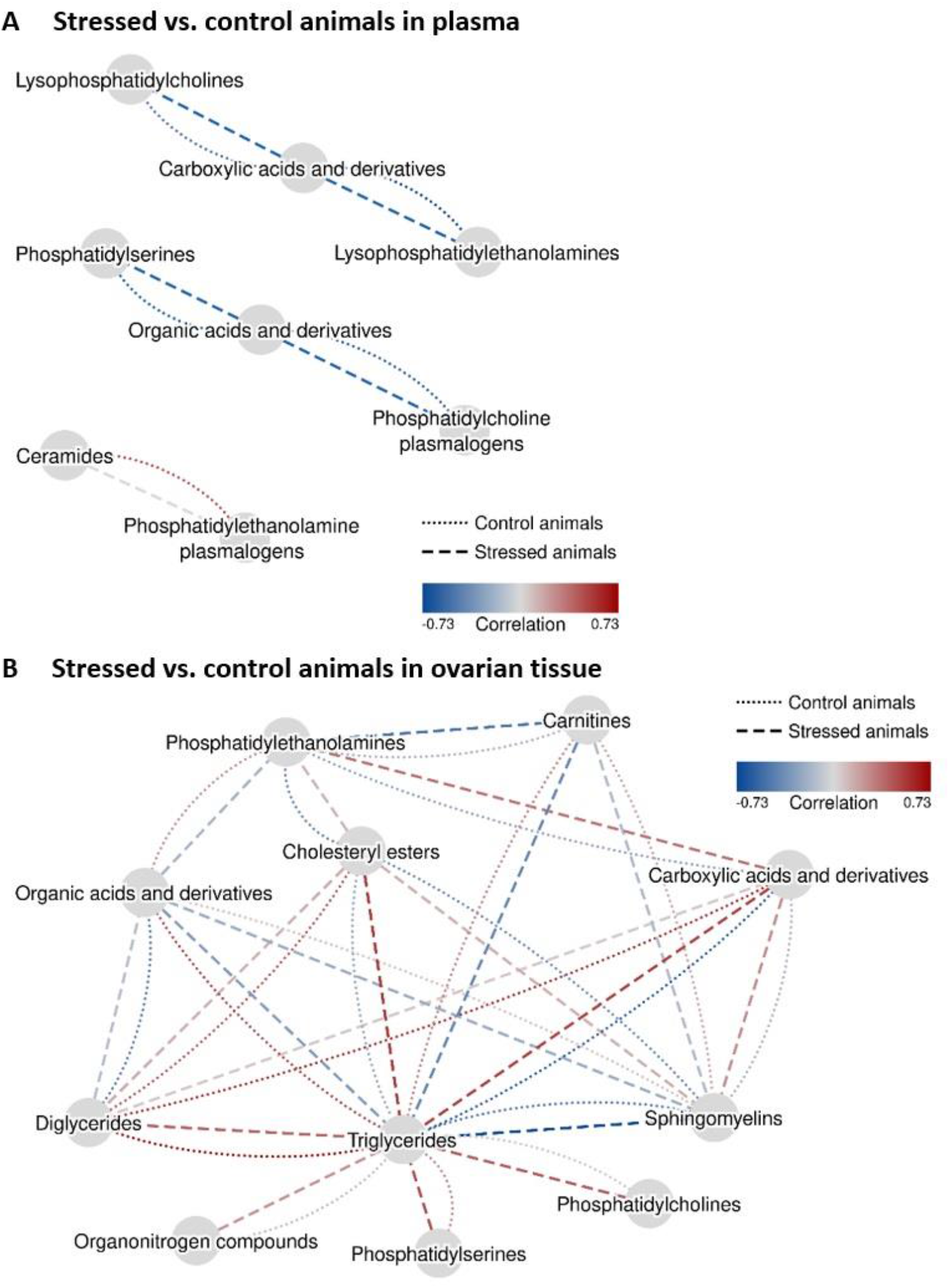
Differential Network Analysis. We conducted differential network analysis (DINGO) to identify a network of statistically significant differences in partial correlations between metabolite classes (summarized using principal component analysis) comparing restraint-stressed mice and controls in plasma (Panel A) and in ovarian tissue (Panel B). The networks show metabolite classes with a statistically significant difference (q-value<0.2) in partial correlations between control and restraint-stressed animals. Each metabolite class is shown as a grey circle. Each pair of metabolite classes is linked by two edges, one representing partial correlations among control animals (dotted line) and one among restrained-stressed animals (dashed line). Positive partial correlations are represented by shades of red while inverse partial correlations are represented by shades blue of blue. Correlations close to zero are shown in shades of grey.

### Metabolites associated with restraint stress in ovarian tissue

The pairwise correlation structure among metabolites measured in ovarian tissue also showed differences between restraint-stressed mice and controls (Figure 1, Panels C and D). Comparing restraint-stressed mice to control mice, 150 metabolites were statistically significantly different at q-value≤0.2 and 30 metabolites at q-value≤0.05 (Figure 2C, Supplementary Table 4). Differences occurred in both directions with some metabolites lower and some higher in restraint-stressed versus control mice. The top five metabolites with the lowest q-values (q-value≤0.05) were DAGs and TAGs, which were consistently elevated in ovarian tissue of restraint-stressed compared to control mice (ranging from 47% for C32:2 DAG to 135% for C46:1 TAG). By contrast, lower levels of several amino acids (5-aminolevulinic acid, putrescine, hydroxyproline, citrulline, carnosine and pyroglutamic acid) were observed comparing restraint-stressed mice versus controls (q-value≤0.1). Considering the magnitude of differences, the largest difference among metabolites with lower levels among restraint-stressed mice versus controls was observed for C20:3 cholesteryl ester (CE) (45% lower; q-value=0.1). Among metabolites with higher levels among restraint-stressed mice versus controls, the largest difference was observed for C44:1 TAG (191% higher; q-value=0.05). Six metabolite classes were enriched at p-adjusted<0.2 (Figure 2D, Supplementary Table 5), including higher levels of TAGs and DAGs (p-adjusted<0.03) and lower levels of CEs, carnitines and PCs and PC plasmalogens (p-adjusted≤0.15) in the restrained-stressed mice.

The ovarian tissue network reflecting differences in the partial correlation between restrained-stressed and control mice included eleven metabolite classes with 20 significant edges (q-value<0.2; Figure 3B, Supplementary Table 6). In this network, TAGs were the most connected metabolite class (N=9), followed by sphingomyelins (SM, N=5). Carboxylic acids and derivatives, CEs, DAGs, organic acids and derivatives, and PEs were each connected to four other metabolite classes while carnitines were connected to three other classes. Organonitrogen compounds, PCs and PSs were each connected to one metabolite class. The largest differences in partial correlations comparing control to restraint-stressed animals were observed between TAGs and carboxylic acids and derivatives (controls: −0.60, stressed: 0.52; q-value<0.01), TAGs and CEs (controls: 0.23, stressed: 0.54; q-value<0.01), TAGs and organic acids and derivatives (controls: 0.50, stressed: −0.3; q-value<0.01), and TAGs and carnitines (controls: 0.27, stressed: −0.41; q-value<0.01).

### Comparison of metabolites associated with restraint stress in plasma versus ovarian tissue

A total of 46 metabolites were significantly associated with restraint stress (q-value<0.2) in both plasma and ovarian tissue (Table 1). Of these, 21 metabolites were consistently inversely associated with restraint stress in both plasma and ovarian tissue, and these metabolites primarily included CEs, PC/PE plasmalogens, and amino acids (putrescine, dimethylglycine, hydroxyproline, citrulline, carnosine, and lysine). The associations of restraint stress with the remaining 25 metabolites were related in the opposite directions within plasma versus ovarian tissues (24 were TAGs and DAGs). All TAGs and DAGs were inversely associated with restraint stress in plasma and positively associated in ovarian tissues. The opposite was observed for C18 carnitine, which was higher in plasma and lower in ovarian tissues comparing restraint-stressed mice versus controls.

**Table 1:** Metabolites significantly associated with stress in plasma or in ovarian tissue. Metabolite with q-value≤0.2 are included in the table. We calculated the percent difference between stressed and control mice as the difference between mean metabolite value among stressed mice and mean metabolite values among control mice divided by the mean metabolite value among control mice.

| Metabolite | HMDBID | Plasma |  |  | Ovarian Tissue |  |  |
| --- | --- | --- | --- | --- | --- | --- | --- |
|  |  | Percent Difference | P-value | Q-value | Percent Difference | P-value | Q-value |
| putrescine | HMDB01414 | -46.14 | 0.001 | 0.01 | -17.59 | 0.01 | 0.05 |
| carnosine | HMDB00033 | -59.69 | 0.001 | 0.01 | -35.16 | 0.043 | 0.1 |
| C36:3 PC | HMDB08105 | -23.93 | 0.003 | 0.034 | -11.98 | 0.065 | 0.11 |
| choline | HMDB00097 | -22.19 | 0.008 | 0.051 | -9.32 | 0.095 | 0.13 |
| C20:4 CE | HMDB06726 | -16.56 | 0.013 | 0.072 | -42.92 | 0.013 | 0.06 |
| C34:0 PI | HMDB09805 | -27.71 | 0.013 | 0.072 | -25.03 | 0.004 | 0.05 |
| citrulline | HMDB00904 | -26.62 | 0.022 | 0.097 | -23.41 | 0.028 | 0.08 |
| C38:6 PC plasmalogen | HMDB11319 | -18.57 | 0.035 | 0.116 | -13.31 | 0.182 | 0.19 |
| C18:1 CE | HMDB00918 | -8.39 | 0.043 | 0.124 | -35.77 | 0.035 | 0.09 |
| C18:2 CE | HMDB00610 | -9.42 | 0.043 | 0.124 | -43.73 | 0.004 | 0.05 |
| dimethylglycine | HMDB00092 | -22.70 | 0.043 | 0.124 | -25.25 | 0.004 | 0.05 |
| 2-deoxycytidine | HMDB00014 | -23.71 | 0.043 | 0.124 | -23.36 | 0.065 | 0.11 |
| hydroxyproline | na | -20.17 | 0.053 | 0.139 | -24.05 | 0.01 | 0.05 |
| C32:0 PC | HMDB07871 | -28.66 | 0.053 | 0.139 | -14.43 | 0.133 | 0.17 |
| C38:6 PC | HMDB07991 | -11.15 | 0.065 | 0.161 | -22.57 | 0.043 | 0.1 |
| C34:1 PC plasmalogen-A | HMDB11208 | -21.85 | 0.065 | 0.161 | -17.36 | 0.065 | 0.11 |
| 5-aminolevulinic acid | HMDB01149 | -19.36 | 0.079 | 0.182 | -23.73 | 0.008 | 0.05 |
| phosphoethanolamine | HMDB00224 | -24.51 | 0.095 | 0.196 | -12.04 | 0.113 | 0.15 |
| C24:0 Ceramide (d18:1) | HMDB04956 | -32.66 | 0.095 | 0.196 | -9.60 | 0.156 | 0.17 |
| lysine | HMDB00182 | -35.42 | 0.095 | 0.196 | -11.71 | 0.156 | 0.17 |
| C38:6 PE plasmalogen | HMDB11387 | -37.88 | 0.095 | 0.196 | -11.99 | 0.156 | 0.17 |
| C58:8 TAG | HMDB05413 | -46.93 | <0.001 | 0.003 | 36.61 | 0.065 | 0.11 |
| C58:9 TAG | HMDB05463 | -50.87 | 0.001 | 0.01 | 59.71 | 0.053 | 0.11 |
| C56:6 TAG | HMDB05456 | -44.83 | 0.002 | 0.027 | 35.99 | 0.065 | 0.11 |
| C54:4 TAG | HMDB05370 | -46.85 | 0.004 | 0.039 | 65.75 | 0.022 | 0.07 |
| C54:5 TAG | HMDB05385 | -44.20 | 0.006 | 0.046 | 59.22 | 0.095 | 0.13 |
| C58:10 TAG | HMDB05476 | -46.02 | 0.008 | 0.051 | 63.98 | 0.156 | 0.17 |
| C58:7 TAG | HMDB05471 | -48.70 | 0.01 | 0.065 | 27.58 | 0.156 | 0.17 |
| C54:3 TAG | HMDB05405 | -43.69 | 0.013 | 0.072 | 65.94 | 0.028 | 0.08 |
| C18 carnitine | HMDB00848 | 42.24 | 0.022 | 0.097 | -19.91 | 0.156 | 0.17 |
| C48:0 TAG | HMDB05356 | -21.90 | 0.028 | 0.107 | 78.26 | 0.001 | 0.05 |
| C36:2 DAG | HMDB07218 | -36.84 | 0.028 | 0.107 | 33.41 | 0.133 | 0.17 |
| C52:2 TAG | HMDB05369 | -31.41 | 0.035 | 0.116 | 62.20 | 0.017 | 0.06 |
| C52:3 TAG | HMDB05384 | -34.54 | 0.035 | 0.116 | 60.98 | 0.01 | 0.05 |
| C54:8 TAG | HMDB10518 | -40.06 | 0.035 | 0.116 | 127.73 | 0.022 | 0.07 |
| C50:2 TAG | HMDB05377 | -29.70 | 0.043 | 0.124 | 66.23 | 0.017 | 0.06 |
| C52:4 TAG | HMDB05363 | -34.40 | 0.043 | 0.124 | 67.66 | 0.053 | 0.11 |
| C56:7 TAG | HMDB05462 | -35.13 | 0.043 | 0.124 | 42.72 | 0.095 | 0.13 |
| C36:3 DAG | HMDB07219 | -37.54 | 0.043 | 0.124 | 31.84 | 0.182 | 0.19 |
| C56:9 TAG | HMDB05448 | -39.67 | 0.043 | 0.124 | 105.88 | 0.022 | 0.07 |
| C56:8 TAG | HMDB05392 | -34.35 | 0.053 | 0.139 | 65.26 | 0.053 | 0.11 |
| C50:4 TAG | HMDB05435 | -45.29 | 0.053 | 0.139 | 111.76 | 0.01 | 0.05 |
| C54:6 TAG | HMDB05391 | -26.93 | 0.079 | 0.182 | 44.36 | 0.035 | 0.09 |
| C52:6 TAG | HMDB05436 | -45.23 | 0.079 | 0.182 | 122.42 | 0.013 | 0.06 |
| C52:7 TAG | HMDB10517 | -61.47 | 0.08 | 0.182 | 149.12 | 0.01 | 0.05 |
| C50:3 TAG | HMDB05433 | -29.31 | 0.095 | 0.196 | 93.16 | 0.017 | 0.06 |

HMDB ID: Metabolite ID in the Human Metabolome Database; PC: phosphatidylcholine; CE: cholesteryl ester; PI: Phosphatidylinositol; TAG: triacylglycerol; na: not available.

## Discussion

To our knowledge, this is the first study to evaluate metabolomic changes due to chronic stress in both plasma and ovarian tissue from female mice. In plasma, the individual metabolites most strongly associated with restraint stress versus control were several LPCs, and the top metabolite classes were carnitines (positively associated), DAGs and TAGs (inversely associated). In ovarian tissue, numerous individual DAGs and TAGs, as well as these metabolite classes as a whole, were increased in mice experiencing restraint stress versus controls, the opposite of that observed in plasma. Similarly, carnitines were under-expressed in stressed mice as a metabolite class in ovarian tissue in contrast to what we observed in plasma. However, several metabolites (CEs, PC/PE plasmalogens and amino acids) were consistently inversely associated with restraint stress in both plasma and ovarian tissue. On the differential network level, fewer connections between metabolite classes were observed in plasma compared to ovarian tissue, suggesting a more severely disturbed metabolic network in response to restraint stress in the local ovarian environment than in circulation.

Prior research in humans and animals across different models of chronic stress has demonstrated that stress is associated with lipid dysregulation. Three animal studies, including one with female mice, reported higher total cholesterol levels in stressed versus non-stressed animals, although results for LDL and VLDL were less consistent (4, 10, 12, 16-19). In human studies, depression has been related to higher circulating levels of TAGs, DAGs, and LPEs (26, 27), whereas lower circulating levels of PCs were observed among individuals with a Type D personality versus other personality types (21). Further, carnitine deficiency has been observed among MDD patients (30), and supplementation of acetyl-L-carnitine had potential benefits on improving depressive symptoms (50, 51). In the current study, we also observed differences in these lipid metabolite classes following exposure to restraint stress, although TAGs, DAGs, and carnitines were inversely associated with restraint stress in plasma. The opposite direction in lipidomic changes in humans and mice is consistent with species-specific stress-induced changes in overall adiposity, given the strong correlations of TAGs, DAGs and carnitines with adiposity. While humans who experience chronic stress tend to gain more weight over time compared with those who do not (52, 53), exposure to restraint stress results in no appreciable difference in body weight in mice (38). It is also possible that differential changes across species may be related to behavioral factors, such as eating patterns, rather than stress per se. Additional prospective assessments in human populations are needed to further assess the apparent lack of conservation between species with respect to stress and lipid dysregulation. Further, in our study, TAGs and DAGs were increased in ovarian tissue versus decreased in plasma in restraint-stressed versus control mice, while the opposite was observed for carnitines. This suggests that chronic stress may lead to the uptake or release of certain lipids between tissue and circulation, although tracing studies would be necessary to confirm this. Overall, the preponderance of the evidence supports that lipid metabolism is dysregulated under conditions of chronic stress, although the changes may differ by species and by tissue compartment.

A novel finding in our study was that levels of several amino acids, such as carnosine, putrescine, dimethylglycine, hydroxyproline, citrulline, and lysine, were reduced under chronic stress in both plasma and ovarian tissue. Putrescine is a polyamine found in nearly all living organisms and is highly conserved, with key roles in transcription, translation, and response to stress, including osmotic and heat stress (54). In Chinese hamster ovarian (CHO) cells, putrescine bound to human beta-adrenoreceptors, a key receptor for catecholamines; the addition of putrescine led to increased cAMP levels, morphological changes in CHO cells, and increased cell migration that was blocked by the addition of propranolol (a beta-blocker) (55). Notably, ovarian tumor tissue had higher levels of putrescine than adjacent normal tissue, and putrescine can alter neoplastic growth (56). These data suggest that stress-related alterations in putrescine may have the potential to influence ovarian carcinogenesis directly (57). Further, there is some evidence suggesting decreased levels of dimethylglycine and citrulline in animals exposed to chronic unpredictable mild stress or in MDD patients (11, 29), whereas supplementation of lysine has been shown to mitigate stress and reduce stress-induced anxiety in individuals with insufficient dietary lysine intake (58). Despite the plausibility of these observations, existing evidence on the relationships between chronic stress and amino acid profiles has been limited and inconsistent. Additional research, using a consistent assay platform across multiple models of stress, is needed.

Findings from the differential network indicate more metabolic perturbations induced by restraint stress in ovarian tissue than in plasma. Comparing ovarian tissue with plasma, there were not only more differential edges, but also more qualitatively different correlations (i.e., in different directions) between metabolite classes. For example, of the four significant differential edges in plasma, all correlations in restraint-stressed mice or controls had the same direction just differing in the strength of the relationship. By contrast, of the 20 significant edges in ovarian tissue, 12 had differential correlations in the opposite direction between restraint-stressed mice versus controls. Given that metabolomics reflect upstream changes at the genomic, transcriptional, and posttranslational levels, our results suggest that the ovaries may be highly susceptible to the influences of chronic stress. This work provides insight into possible metabolic dysfunction that may promote the ovarian cancer growth and progression. Our findings also provide basis for future human studies to explore local metabolomic changes at tissue-specific levels in response to stress and their relationships with cancer risk.

This study has several strengths, including assessing female mice using a well characterized chronic stress system, leveraging a well-validated metabolomics platform, and simultaneously considering both plasma and ovarian tissue. Although the sample size was relatively small and some associations were likely missed, we still were able to identify the most significant metabolomic changes related to chronic stress even after correcting for testing multiple correlated hypotheses. One challenge in comparing our results with prior studies is that different metabolomics assay platforms do not assess the same metabolites. For example, some studies (including ours) measured lipids by the number of carbons and double bonds, and by backbone type (e.g., C22:6 LPC), while others measured specific fatty acids and related salts (e.g., octadecanoic acid). Efforts to standardize platforms across multiple studies are critical to enhance comparability and allow for direct validation across studies.

In summary, we identified differences in multiple lipid and amino acid metabolites in plasma and ovarian tissue of female mice after exposure to restraint stress. These metabolites included TAGs, DAGs, carnitines, putrescine, dimethylglycine, hydroxyproline, citrulline, and lysine, among others. Importantly, some, but not all, of the affected metabolites (primarily TAGs and DAGs) occurred primarily either in plasma (a marker of systemic influences) or in ovarian tissue (representing local changes), suggesting that research to understand the biological impact of chronic stress needs to consider both systemic and tissue-specific alterations. Replication of our findings in larger studies considering different methods/forms of inducing chronic stress, in mice and human populations, are needed, and clinical implications of the stress-related metabolomic profiles remain to be elucidated.

## Acknowledgments

This work was supported, in part, by NIH grants R01 CA163451, P50CA217685, R35CA209904, and P30CA016672; the Ovarian Cancer Research Alliance; the American Cancer Society Research Professor Award; and the Frank McGraw Memorial Chair in Cancer Research. T.H. was supported by grant K01HL143034 from the National Heart, Lung and Blood Institute. C.J.P. was funded by NIEHS R01ES032470. K.H.S, L.D.K. and S.E.H were supported by grant R01AG051600 from the National Institute on Aging.

